# Subconcussive head impacts sustained during American football alter gut microbiome diversity and composition

**DOI:** 10.1101/2024.07.05.602235

**Authors:** Zack Pelland, Aziz Zafar, Ahmet Ay, Ken Belanger

**Author notes:** These authors contributed equally to this project (Zack Pelland, Aziz Zafar).

## Abstract

Subconcussive head impacts (SHIs) are a public health concern amongst at-risk populations. SHIs are hits to the head that do not typically generate symptoms and are unlikely to meet diagnostic criteria for mild traumatic brain injury (mTBI). Changes in the gut microbiome have been associated with mTBIs and implicated in both acute recovery from and neurodegenerative pathologies associated with repeated mTBI. This study monitored the gut microbiomes and head impact exposure of collegiate American football players across a competition season to determine if SHIs lead to acute and longer-term changes to the gut microbiome. SHI exposure correlates with changes in microbial diversity and composition three days post-exposure, the athletes’ gut microbiomes change significantly across the season, and mixed effects modeling provides evidence for cumulative effects of SHIs. These data provide the first evidence for a link between SHIs and changes in the diversity and composition of the gut microbiome.

## INTRODUCTION

An estimated 47 million people across the globe suffer from a mild traumatic brain injury (mTBI) every year^1–3^. mTBIs are acute injuries to the brain caused by external physical forces applied to the head or secondary forces applied to other parts of the body, commonly resulting in confusion, a brief loss of consciousness, amnesia, and other transient neurological symptoms^4^. Approximately 1 in 6 mTBI patients experience short-term disabilities due to symptoms prolonging for three or more months following an injury^5,6^, and repeated exposure to mTBIs—which is common amongst military personnel and contact sport athletes^7^—has been linked to chronic disability and an increased risk of developing neurodegenerative and psychiatric diseases^7–15^.

The short-term disabilities and neurodegenerative pathologies that may develop after repeated exposure to mTBIs have been linked to chronic inflammation^16–31^. The gut microbiome, the community of trillions of microorganisms that line the intestinal tract, has been shown to be a key regulator of inflammation and the neuroimmune system^32–35^. Numerous neurological conditions with related neuroinflammatory characteristics have been associated with abnormalities in the composition of the gut microbiome, known as gut dysbiosis^36–39^. Recent findings suggest that mTBIs similarly cause gut dysbiosis^40–50^, which, due to the gut microbiome’s role in regulating inflammation, may play a role in the short-term disability and the development of neurodegeneration linked to repetitive mTBIs^51,52,52,53^. Identifying and treating gut dysbiosis has shown the potential to diagnose and alleviate the symptoms of other neurological and psychiatric conditions^39,54,55^. Thus, recognizing and characterizing alterations in the gut microbiome following mTBIs represents a potential avenue to predict, treat, and prevent severe or long-term neurological damage from mTBIs^50,56–58^.

While mTBIs have been more substantially explored, subconcussive head impacts (SHIs) have been less so given the challenges in monitoring these injuries. SHIs consist of cranial impacts or rapid accelerations of the body that likely contribute to minor brain injuries but are not recognized or clinically diagnosed as an mTBI^59^. Repeated exposure to SHIs is particularly common among American football players, with athletes experiencing between 100 and 1,000 head impacts across a season^59–61^. Similar to mTBIs, the accumulation of SHIs has been linked (to a lesser extent) to acute changes in inflammatory markers, short-term changes in cognitive function, and an increased risk of neurodegeneration or early cognitive decline^59,61–69^. However, to our knowledge, the link between SHIs and gut microbiome composition has not yet been investigated.

Given the connections between the gut microbiome and mTBIs and the frequent exposure to SHIs among American football players, we tested two hypotheses: whether individual or accumulated impacts on a single day lead to acute changes in the gut microbiome, and whether SHIs have a cumulative effect that leads to longer-term changes in the gut microbiome. To test these hypotheses, we monitored the gut microbiomes of National Collegiate Athletic Association (NCAA) Division I American football players across a competition season while tracking head impacts sustained during all practices and games. We found that microbial diversity in athletes changed within three days after substantial subconcussive head impact exposure. Specifically, we observed correlations between changes in the abundance of Coriobacteriales, *Prevotella*, and *Ruminococcus* and head impact load sustained in the past 48 to 72 hours. We also found that the athletes’ gut microbiomes change significantly across the season, with evidence that this change is partially due to the cumulative effects of SHIs and other clinical factors. These data provide the first evidence for a link between SHIs and acute and long-term changes in the gut microbiome.

## METHODS

### General study design

Nineteen NCAA Division I American football team members had their head impacts, on-field physical activity, relevant clinical factors, and fecal microbiome monitored across a competition season, beginning during pre-season training (Figure 1). After exclusions, 226 fecal samples from the six remaining participants were analyzed. Head impacts were monitored using the Riddell_®_ InSite helmet-based impact monitoring system (Riddell_®_), on-field physical activity with portable 10 Hz GPS units (Catapult S7 and G7, Catapult Sports), clinical factors through digital surveys, and the fecal microbiome characterized with 16S rRNA PCR amplification and next-generation sequencing.

**Figure 1.**
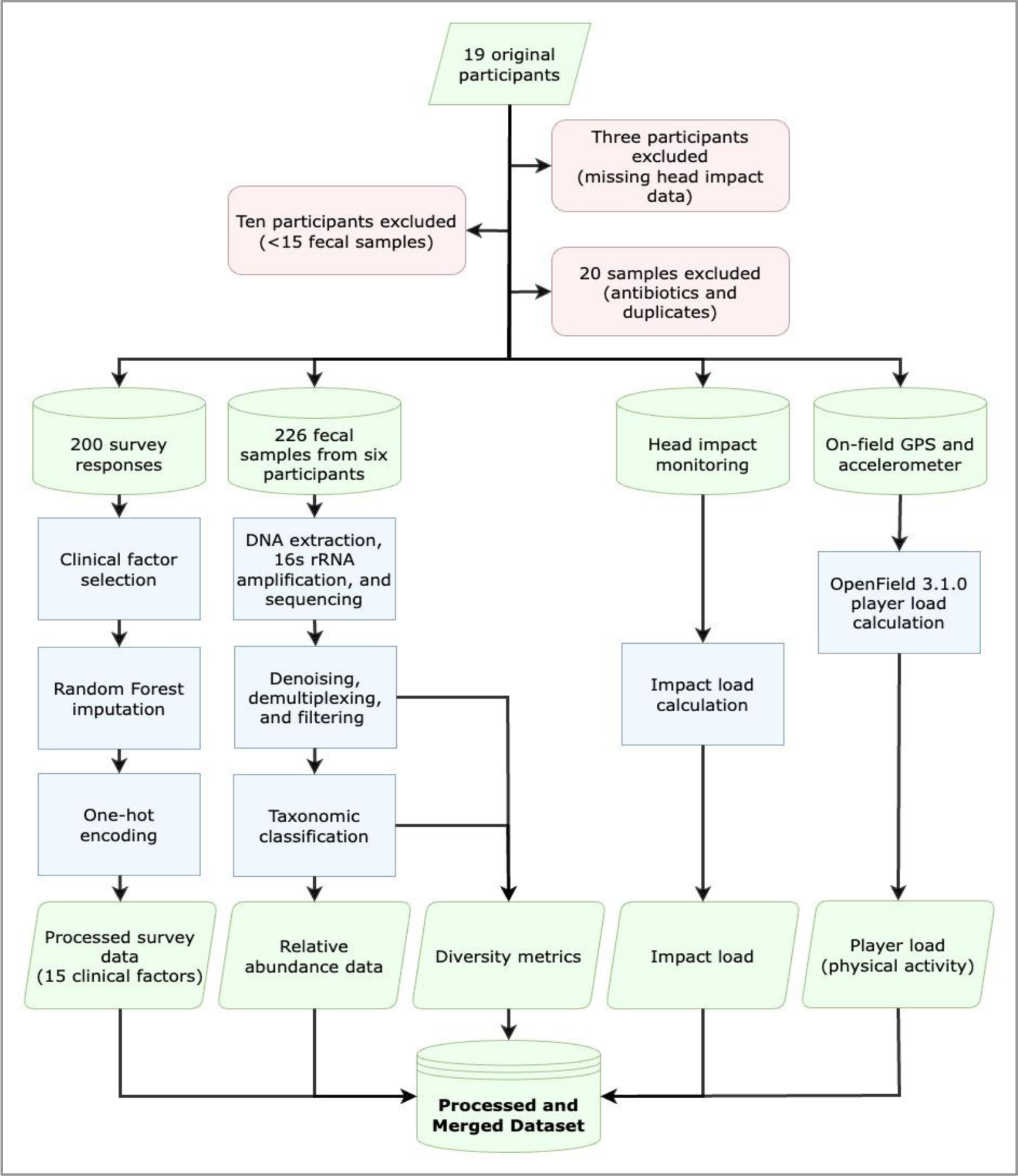
Data collection and processing flowchart. Head impact data, on-field activity data, surveys of clinically relevant factors, and fecal samples were collected from 19 original participants. Thirteen participants were excluded due to missing head impact data or insufficient fecal samples (fewer than 15 samples). Twenty fecal samples from the remaining six participants were excluded due to oral antibiotic use and duplicate samples collected on the same day. Fifteen clinical factors from available survey data were selected, and missing survey data was imputed. The DNA from 226 fecal samples was extracted, sequenced, and processed into relative abundance data and diversity metrics. Head impacts were monitored, and impact load was calculated based on the severity of impacts. On-field activity was monitored to measure physical strain (player load).

### Participants

This single-site prospective cohort study recruited 19 active male NCAA Division I Football Championship Subdivision football team members through a survey sent to the entire team (∼90 players). Participants were excluded from the study if they had been prescribed oral antibiotics or had sustained a concussion in the past 60 days. Three subjects of the initial 19 were excluded from analysis due to all or part of their head impact data not being collected, and ten more subjects were excluded from analysis due to collecting fewer than 15 fecal samples. Only the data for the six remaining participants was analyzed. Prior to the commencement of sample collection, participants reported medical history, sports history, and demographic information (see Supplementary Document 1 for the survey). Participant positions are not provided to protect participant confidentiality. All study procedures were approved by the Institutional Review Board of Colgate University and performed in accordant with relevant guidelines and regulations. Informed consent was obtained from all participants prior to data collection.

### Head impact data

Head impact data was collected using the Riddell_®_ InSite helmet-based impact monitoring system (Riddell_®_). The Riddell_®_ InSite system includes player units and helmet liners that track and characterize impacts to the helmet based on the linear force at one of five levels (15-19 g, 20-28 g, 29-43 g, 44-63 g, and 63+ g) and location in one of five regions of the helmet (front, crown, back, left, and right). In the present study, an impact load score was calculated by granting head impacts between 15-19 g a score of 17, 20-28 g a score of 24, 29-43 a score of 36, 44-63 g a score of 53.5, and 63+ g a score of 63. The sum of these scores for head impacts sustained during each practice and game was calculated to generate a head impact load score for each session. When an impact load score of 70 or above was sustained for any practice or game session, participants were defined as having experienced “substantial head impact exposure.” Practice and game sessions were designated a specific time point to determine when head impacts were sustained in reference to fecal sample collections; the time point for each head impact load score was designated to be the midpoint of the practice or game (i.e., head impacts sustained during a practice session from 4:00 PM to 6:00 PM were designated to have occurred at 5:00 PM). Head impact location was not taken into consideration for this study.

### On-field activity monitoring

The activity profiles of four of the six participants and 14 total team members were monitored during every training and competition session with portable 10 Hz GPS units (Catapult S7 and G7, Catapult Sports). The GPS unit was worn in a vest and rested between the shoulder blades, which did not limit upper limb or torso movements. In addition to the GPS-derived velocity, distance, and acceleration values, the GPS units also provided inertial movement analysis (IMA) data based on integrating 100 Hz microsensor data (accelerometer, gyroscope, and magnetometer). The data was processed by dedicated software (Catapult OpenField Version 3.1.0) to generate the player load metric, which was exported for analysis. Participants 8 and 16 did not wear designated activity monitors, and the team average player load (calculated from all 14 players) was assigned to these participants for each practice and game session.

### Fecal sample collection lifestyle questionnaires

Participants were asked to complete a brief questionnaire following each fecal sample collection as well as more comprehensive background surveys before and after the season. The daily questionnaires (Supplementary Files 1 and 2) assessed clinical factors pertaining to the experiences and behaviors of the participants during the 24 hours before each fecal sample collection. Some clinical factors were excluded from the analysis due to low or inaccurate reporting. Fifteen clinical factors were kept in the final analysis, including a stress rating modified from the PSS-10^70^, a selection of SCAT5^71^ symptoms, sleep quantity, sleep quality, illness, vomiting, caffeine use, NSAID use, supplement use, alcohol use, nicotine use, Bristol Stool Form Scale^72^, orthopedic injury, and other potentially relevant factors (Supplementary Table 1). Participants occasionally did not complete the questionnaire after a sample collection. For fecal samples not accompanied by a questionnaire, the survey data was imputed using the miceRanger library in R. When the time of day of fecal sample collection was not recorded, the participant’s average time of day of collection was used.

### Fecal sample collection and storage

Participants were instructed to collect fecal samples from daily bowel movements using the Norgen Biotek Corp Stool Collection Kit (Norgen Biotek Corp) or Protocult^TM^ Collection Device (Therapak, LLC). All fecal samples were submerged in Norgen Biotek nucleic acid preservative (Norgen Biotek Corp) and stored at -20°C immediately after collection. 246 fecal samples were collected from the remaining six participants throughout the study. Fifteen fecal samples from participant 9 and four samples from participant 16 were excluded from the analysis after they were prescribed oral antibiotics. Participant 9 collected two fecal samples on one day, and the average taxonomic and diversity data was taken for these samples, leaving 226 total fecal samples for analysis.

### DNA extraction and sequencing

DNA was extracted from the fecal samples using the DNeasy PowerSoil Kit (Qiagen) according to the manufacturer’s instructions. 16S rRNA PCR amplification and next-generation sequencing were performed at MR DNA (www.mrdnalab.com) using primers 515F-Y (5’-GTGCCAGCMGCCGCGGTAA-3’)^73^ and 806R (5’-GGACTACHVGGTWTCTAAT-3’)^74^ using Illumina MiSeq (Illumina Corp) 2x300 paired-end reads. Control samples were sequenced, which verified negligible contamination. These included an internal mock (ATCC), a negative control that went through the extraction process, a negative control consisting of stock solution, and two positive controls consisting of *Shigella spp.* (Supplementary Figure 1). We obtained 14,276,903 16S rRNA V4 region sequences over the 226 samples sequenced, providing an average of 58,036.19 (SD 18,032.83) reads per sample. The minimum forward and backward trimmed sequence lengths used in the analysis were 220 and 180, respectively.

### Data processing - Taxonomic data

For each sample, forward and reverse reads were processed using QIIME2’s processing pipeline^75^. The DADA2 method was used for denoising, with 220 base pairs truncated for the forward read and 180 for the reverse read. The truncation length was identified using visual assessment, considering a trade-off between information and the data size. For the obtained amplicon sequence variants (ASVs), any features that appeared in less than three samples or had absolute abundance below ten were removed. ASVs were collapsed into taxonomic levels using the Silva V4 Classifier.^76^ The classifier provided an abundance table for Operational Taxonomic Units (OTUs), such as specific microbial orders or species. Relative abundances of OTUs were computed and used for each sample for further analysis. Due to our limited sample size, analysis was done on the eight most abundant orders and twenty-one specific taxa (1 order, 11 families, 8 genera, and 1 species) reported to be significantly related to brain injuries in the literature and available in the present dataset (Supplementary File 3)^45–48,50,77–79^.

### Data processing - Alpha and beta diversity

QIIME2 was used to compute Faith’s Phylogenetic Diversity using ASVs directly with an RAxML phylogenetic tree and Simpson’s Diversity using species-level abundances for each stool sample to measure alpha diversity within each sample^80^. Faith’s Phylogenetic Diversity is calculated by first generating a phylogenetic tree using similarities between sequences^81^. Beta diversity was calculated for each sample collected with respect to that individual’s baseline sample collected before the first preseason practice session (Day 1) using Bray-Curtis Dissimilarity measurement^78^.

### Data analysis - Data slicing and repeated-measures analysis

To understand the association between head impact and diversity metrics over time, we identified practice and game sessions in our data where a player sustained a head impact load greater than or equal to the 75th percentile of all head impacts and did not sustain any head impacts of this magnitude for the following three days. The cutoff for the 75th percentile of head impact exposure was a day in which an impact load score of 70 was sustained; participants were defined as having experienced “substantial head impact exposure” when they sustained an impact load score above this threshold for any practice or game session (Supplementary Figure 2). Changes across a season were analyzed by taking the first three, the center three, and the last three data points available for each player, excluding postseason samples (Supplementary Figure 2). Assumptions of repeated-measures ANOVA, such as Mauchly’s sphericity and assumptions of normality, were not always satisfied. Therefore, Friedman’s Chi Square Rank Test was consistently used for repeated measures to identify statistically significant differences between the diversity metrics within 72 hours of impact (in increments of 24 hours) and throughout the season, using early, middle, and late as our time points. For significant results from Friedman’s Chi Square Rank Tests, Nemenyi’s post hoc pairwise testing was performed.

### Data analysis - Mixed effects linear models

Mixed effects models were used to find linear relationships between the clinical factors we measured (Supplementary Table 1) and the head impact load, and various microbial metrics, including alpha diversity, beta diversity, and the relative abundances of microbial orders, families, genera, and species. A random intercept was used to account for each player, and models were fitted using the lmerTest package^83^ in R (version 4.3.1) and adjusted for multiple testing using Benjamini-Hochberg adjustment^84^. The same confounding variables were used for each diversity metric and taxon of interest. However, head impact load was analyzed with 24 and 48 hour offsets to identify time-dependent correlations between head impact and changes in the gut microbiome. Charts with the results from all model runs can be seen in Supplementary File 4.

### Data analysis - Principal coordinate analysis

To identify and visualize general patterns in the samples collected across the season, a distance matrix was created using Bray-Curtis dissimilarities using species level information. The principal coordinates of this distance matrix were computed and plotted as the first two axes to visualize clustering by player. A circle was plotted around each player’s data points using the mean of the centroid as the center and the maximum Euclidean distance between two points within the same player as the radius of the circle.

## RESULTS

### Data collection overview, participant characteristics, and head impact exposure

To investigate potential links between head impacts and gut microbiome composition, six NCAA Division I football players collected fecal samples, completed questionnaires after fecal sample collection, had head impacts monitored, and had on-field physical activity monitored across a competition season (Figure 1). All six participants were male, white or Caucasian, 21 or 22 years old, and received most of their meals from the same meal plan (see Supplementary Table 2 for further demographics). Two participants began antibiotic use during the season, and fecal samples collected after initiating antibiotic treatment were not included in our analysis (Figure 2a).

**Figure 2.**
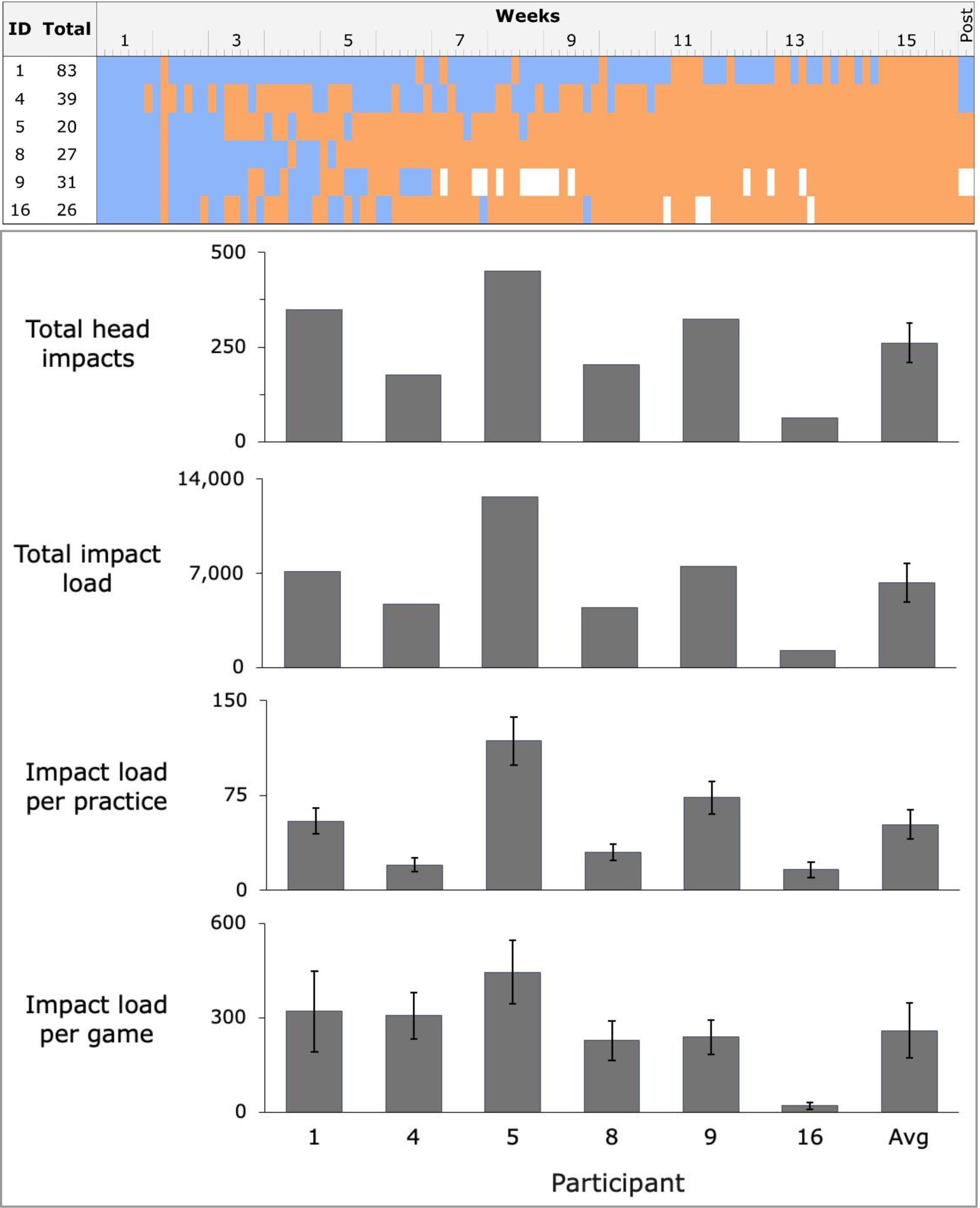
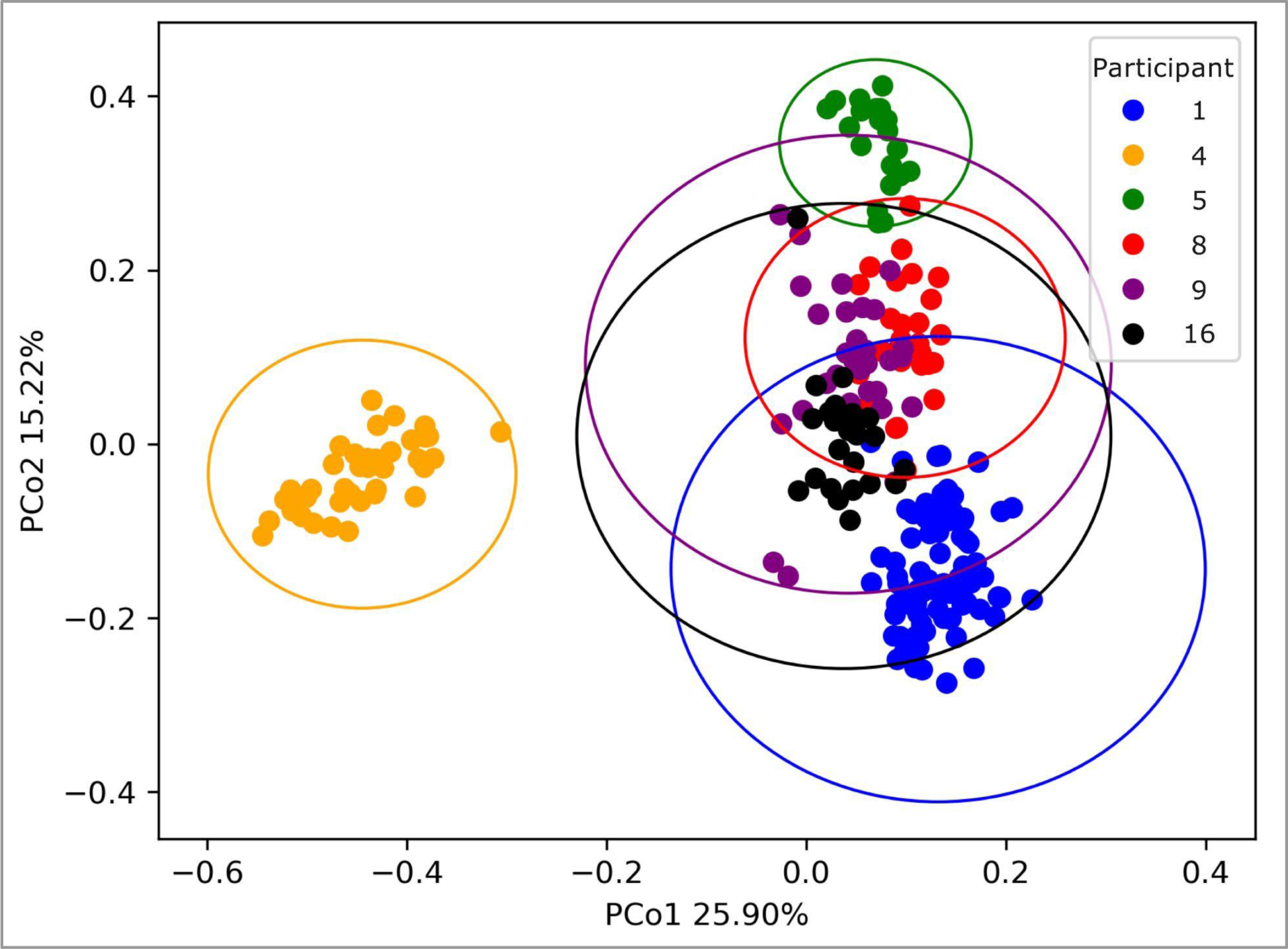
Study participants exhibit variability in head impact frequency and intensity, and in fecal microbiome composition. **(a)** *Availability of fecal samples by participant*. The days on which samples were collected and not collected are marked in blue and orange, respectively. White boxes indicate that the sample collected on this date was excluded due to oral antibiotic use. The final two samples from participants 1, 4, and 9 were collected 10-21 days after the season’s final game. **(b)** *Head impacts and head impact load sustained by participants*. Error bars represent the standard error of the mean. See Methods for data collection and Supplementary Table 2 for exact values. **(c)** *Beta diversity analysis of the gut microbiota from all samples used in analysis*. The scatter plot displays a principal coordinate analysis (PCoA) with data points color-coded by the participant. Each participant’s data is enclosed within its own ellipsoid.

The consistency of fecal sample collection, questionnaire completion, and head impact load exposure varied considerably across the season and among participants (Figure 2a, 2b, Supplementary Table 2, Supplementary Figure 2). Participants sustained an average of 261 head impacts (SEM 51.9) and an average impact load score of 6,308.3 (SEM 1,435.2) across the season, which accumulated from an average of 52.0 (SEM 11.8) impact load sustained per practice session and 260.0 (SEM 86.9) impact load sustained per game session (Figure 2b, Supplementary Figure 2). In our PCoA, we observed a distinct clustering of individual participants.

To examine the relatedness of overall bacterial microbiome composition across the six participants and 226 samples, we performed principal coordinates analysis. The resulting scatter plot (Figure 2c) illustrates the distribution of the dataset along the first two principal coordinates. Notably, participants 4 (orange), 5 (green), and 8 (red) show low-variance clusters, indicating that their microbiome collected at different time points is similar along the first two principal coordinates. In contrast, participants 1 (blue), 9 (purple), and 16 (black) show a broader spread, suggesting higher temporal variability within their microbiomes. We also observed a considerable overlap between some participants, such as participants 8 (red) and 9 (purple), suggesting similarities in their microbiome compositions.

### Bray-Curtis Dissimilarity increases three days following substantial head impact exposure

To examine whether subconcussive head impacts on a single day correlate with changes in gut microbiome composition, we analyzed Bray-Curtis Dissimilarity for each 24-hr period after substantial head impact exposure through 96-hrs post-impact. For this analysis, we only considered samples with an impact load in the top 75th percentile (impact load score of 70 or greater) on day one, followed by 3 consecutive days without any head impacts in the 75th percentile (see methods and Supplementary Figure 2). A total of 13 four-day periods met this criterion, with seven coming from Participant 1, one from Participant 4, two from Participant 5, one from Participant 8, two from Participant 9, and none from Participant 16 (Figure 2). Friedman’s Chi-Squared test on the Bray-Curtis Dissimilarity of the four groups revealed statistically significant differences in bacterial composition across the four days (χ^2^ = 13.02, p = 0.005, df =13.02) (Figure 3). Nemenyi’s post hoc pairwise test revealed that Bray-Curtis Dissimilarity was significantly higher 48-72 hours (0.434 SEM 0.071, Q57 = 2.85, p = 0.025) and 72-96 hours (0.435 SEM 0.096, Q57 = 3.10, p = 0.012) following substantial head impact exposure when compared to 0-24 hours post (0.415 SEM 0.077) (Figure 3).

**Figure 3.**
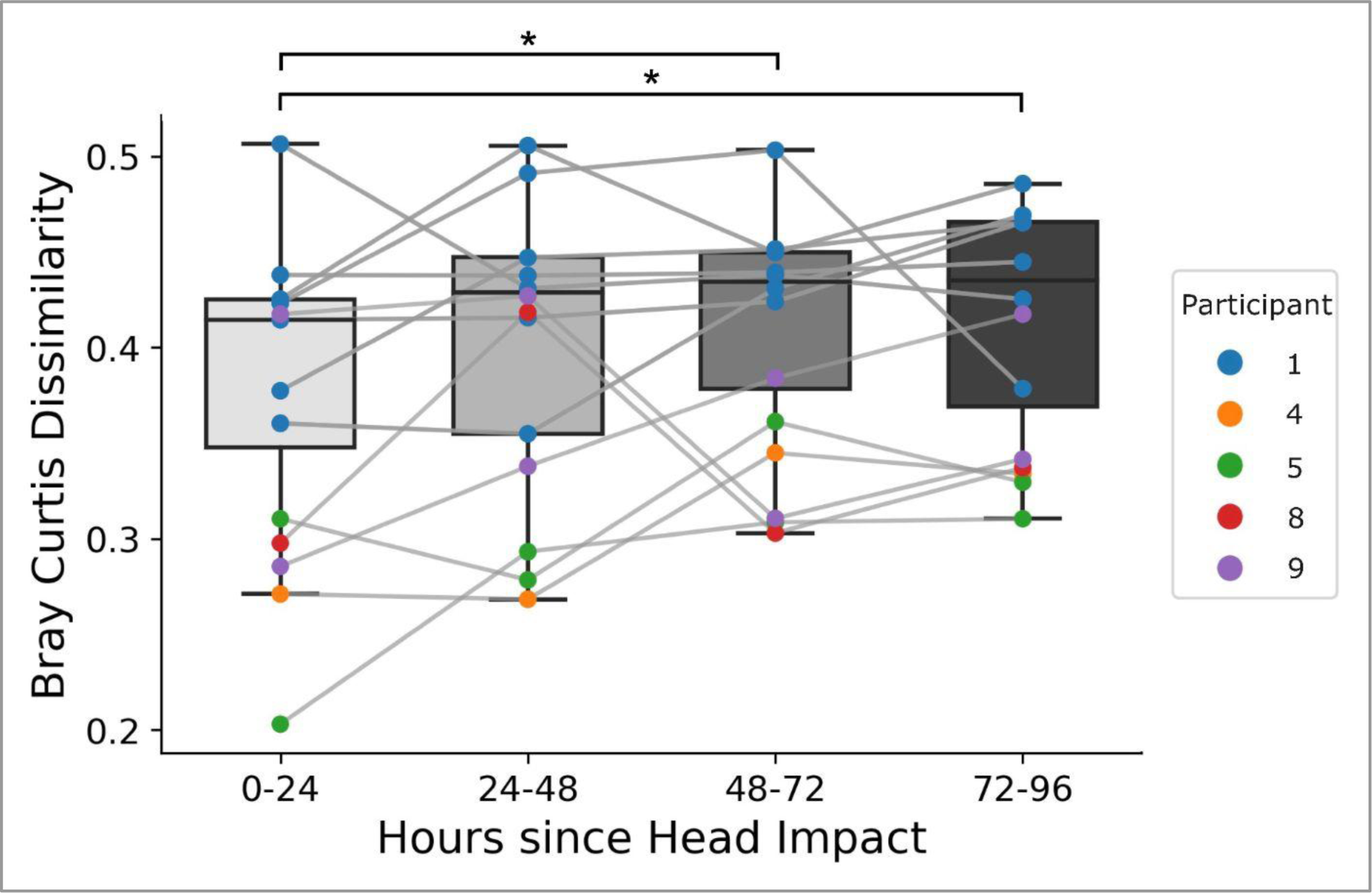
Bray Curtis Dissimilarity increases 48-72 and 72-96 hours following severe head impact exposure. Data slices were isolated such that hour 0 represents a daily head impact load exposure in the top 75^th^ percentile, followed by 96 consecutive hours in which there was no head impact load exposure in the top 75^th^ percentile. Friedman’s Chi-Square test and Nemenyi’s post hoc pairwise test revealed a significant increase in Bray-Curtis Dissimilarity between 48-72 and 72-96 hours post-impact compared to 0-24 (day of impact). Gray lines connect samples from the same data slice. (* indicates p < 0.05)

### Mixed-effect models accounting for confounding variables reveal effects of head impact exposure on microbial diversity

To determine whether head impact load significantly affects Bray-Curtis Dissimilarity, we performed mixed-effect linear modeling that accounted for 15 potentially confounding variables that could influence gut microbiome composition (Supplementary Table 1). We examined changes in microbial composition as time passed from head impact by time-shifting Impact load by 0-24, 24-48, or 48-72 hours under three separate models to determine when an effect on Bray-Curtis Dissimilarity might occur relative to head impact exposure or changes in other recorded variables.

Across all three time-shifted models, time (the progression of time across the entire study period; p < 0.001, t = 3.71, df = 194.96), player load (a measure of physical activity intensity during each practice and game; p < 0.001, t = 3.58, df = 195.25), taking non-steroidal anti-inflammatory drugs (NSAIDs) (p = 0.0194, t = 2.36, df = 195.89), and consuming pre-workout energy drinks (p < 0.001, t = 3.55, df = 195.89) were associated with increases in Bray-Curtis Dissimilarity (Table 1). Type 2 stool character (p = 0.0125, t = -2.52, df = 193.28), type 3 stool (p = 0.0083, t = -2.67, df = 193.79), and stress (p = 0.0201, t = -2.35, df = 169.96) were correlated with decreases in Bray-Curtis Dissimilarity. After adjusting for the false discovery rate using the Benjamini-Hochberg adjustment, only the associations between time and Bray-Curtis Dissimilarity (p = 0.0065), player load (p = 0.0065) and consumption of pre-workout energy drinks remained significant (p = 0.0065).

**Table 1.** Bray-Curtis dissimilarity changes in relation to head impacts sustained in the past 48-72 hours, after adjusting for clinical factors. Significant p-values (< 0.05) are highlighted in dark blue, and marginally significant p-values (< 0.10) are highlighted in light blue. The 48-72 hour time-shifted model is bolded. Only time, player load, and consumption of pre-workout drinks remain significant after the Benjamini-Hochberg p-value correction. “Estimate” indicates the direction and magnitude of the effect on Bray-Curtis Dissimilarity.

| Variable | 0-24 hours post |  | 24-48 hours post |  | 48-72 hours post |  |
| --- | --- | --- | --- | --- | --- | --- |
|  | Estimate | p (adjusted p) | Estimate | p (adjusted p) | Estimate | p (adjusted p) |
| Impact Load | 0.0001538 | 0.1589 (0.3080) | 0.0000334 | 0.7605 (0.8732) | **0.0002870** | **0.0420 (0.1447)** |
| Time (hour) | 0.0000278 | 0.0202 (0.0846) | 0.0000416 | 0.0011 (0.0104) | **0.0000456** | **0.0003 (0.0065)** |
| Impact load*time | -0.0000001 | 0.2291 (0.4177) | 0.0000000 | 0.5217 (0.6738) | **-0.0000002** | **0.1195 (0.2850)** |
| Player load | 0.0001332 | 0.0001 (0.0033) | 0.0001294 | 0.0001 (0.0023) | **0.0001163** | **0.0006 (0.0065)** |
| Orthopedic injury | 0.0106443 | 0.5419 (0.6462) | 0.0093275 | 0.5810 (0.7162) | **0.0103550** | **0.5394 (0.6689)** |
| Type 1 - Severe constipation | -0.0905658 | 0.0218 (0.0846) | -0.0657809 | 0.2540 (0.3937) | **-0.0748907** | **0.1899 (0.2896)** |
| Type 2 - Minor constipation | -0.0581556 | 0.0066 (0.0509) | -0.0564121 | 0.0092 (0.0476) | **-0.0536494** | **0.0125 (0.0646)** |
| Type 3 - Normal | -0.0286114 | 0.0337 (0.0950) | -0.0396518 | 0.0031 (0.0192) | **-0.0353654** | **0.0083 (0.0516)** |
| Type 6 - Minor diarrhea | 0.0039469 | 0.8376 (0.8376) | 0.0272812 | 0.1388 (0.2868) | **0.0252270** | **0.1668 (0.2896)** |
| Type 7 - Severe Diarrhea | 0.0893258 | 0.2563 (0.4210) | 0.0930181 | 0.0576 (0.1984) | **0.0955169** | **0.0515 (0.1597)** |
| Less caffeine | 0.0270608 | 0.2716 (0.4210) | -0.0170889 | 0.5001 (0.6738) | **-0.0134611** | **0.5898 (0.6874)** |
| Normal caffeine | -0.0208674 | 0.2599 (0.4210) | -0.0321247 | 0.1020 (0.2433) | **-0.0254335** | **0.1913 (0.2896)** |
| More caffeine | -0.0325225 | 0.4381 (0.5659) | -0.0967378 | 0.0810 (0.2281) | **-0.0828583** | **0.1321 (0.2896)** |
| Less NSAIDs | 0.0597370 | 0.1012 (0.2413) | 0.0542994 | 0.1308 (0.2868) | **0.0507840** | **0.1516 (0.2896)** |
| Normal NSAIDs | 0.0320098 | 0.0297 (0.0919) | 0.0349986 | 0.0138 (0.0611) | **0.0327675** | **0.0194 (0.0778)** |
| More NSAIDs | -0.0629073 | 0.3335 (0.4700) | 0.0298070 | 0.4582 (0.6738) | **0.0298175** | **0.4525 (0.6100)** |
| Less nicotine | -0.0099746 | 0.6902 (0.7641) | -0.0333357 | 0.2236 (0.3648) | **-0.0353403** | **0.1858 (0.2896)** |
| Normal nicotine | -0.0096307 | 0.6791 (0.7641) | -0.0014977 | 0.9525 (0.9525) | **-0.0017503** | **0.9436 (0.9922)** |
| More nicotine | 0.0502184 | 0.3070 (0.4532) | 0.0691776 | 0.2091 (0.3601) | **0.0773147** | **0.1539 (0.2896)** |
| Pre-workout drink | 0.0617364 | 0.0034 (0.0354) | 0.0859512 | 0.0001 (0.0023) | **0.0762225** | **0.0005 (0.0065)** |
| Number of diet sodas | -0.0036543 | 0.8349 (0.8376) | -0.0032012 | 0.8614 (0.9400) | **-0.0077491** | **0.6696 (0.7414)** |
| Alcohol | 0.0019122 | 0.5417 (0.6462) | 0.0003636 | 0.9025 (0.9400) | **-0.0000051** | **0.9986 (0.9986)** |
| Hours of sleep | 0.0202316 | 0.0184 (0.0846) | 0.0152184 | 0.0790 (0.2281) | **0.0144298** | **0.0908 (0.2559)** |
| Sleep quality | -0.0141333 | 0.1549 (0.3080) | -0.0065948 | 0.4967 (0.6738) | **-0.0050543** | **0.5987 (0.6874)** |
| Stress rating | -0.0082775 | 0.0126 (0.0783) | -0.0075119 | 0.0267 (0.1036) | **-0.0078570** | **0.0201 (0.0778)** |
| Symptom score | -0.0079122 | 0.3503 (0.4722) | -0.0044119 | 0.6007 (0.7162) | **-0.0051181** | **0.5348 (0.6689)** |
| Minor illness | -0.0400530 | 0.0985 (0.2413) | -0.0363080 | 0.1676 (0.3247) | **-0.0293673** | **0.2603 (0.3668)** |
| Mild illness | 0.0782467 | 0.0258 (0.0889) | 0.0468942 | 0.1924 (0.3508) | **0.0458310** | **0.1962 (0.2896)** |
| Vomitted | 0.0120149 | 0.7796 (0.8333) | -0.0049708 | 0.9097 (0.9400) | **0.0021367** | **0.9602 (0.9922)** |
| Change in living or dining | -0.0372360 | 0.1305 (0.2889) | -0.0383404 | 0.0903 (0.2333) | **-0.0353493** | **0.1120 (0.2850)** |

Impact load exposure 48 to 72 hours prior was associated with increased Bray-Curtis Dissimilarity (p = 0.0420, t = 2.05, df = 193.13) (Figure 3, Table 1). After adjusting for the false discovery rate using the Benjamini-Hochberg adjustment, the association between impact load and Bray-Curtis Dissimilarity was no longer significant (p = 0.1447). See Supplementary File 3 for all diversity model runs.

### Gut microbiome composition changes 48-72 hours post head impact

To identify the microbial taxa changes that contribute to changes in Bray-Curtis Dissimilarity 48 to 72 hours following head impact exposure, we performed mixed-effect linear modeling for relevant taxa (see Methods). Decreases in the relative abundances of the order Coriobacteriales, the family *Prevotellaceae,* and the genus *Prevotella* were significantly correlated with higher impact load (p = 0.0154, t = -2.44, df = 192.89; p = 0.0045, t = -2.87, df = 190.21; p = 0.0035, t = -2.95, df = 190.20; respectively) (Table 2). Increases in the relative abundances of the genus *Ruminococcus* and the order *Verrucomicrobiales* were significantly or marginally correlated with higher impact loads (p = 0.0075, t = 2.70, df = 190.68; p = 0.0776, t = 1.78, df = 195.05; respectively) (Table 2). The interaction effect of impact load and time (impact load*time) was significant or close to being significant for all taxa mentioned (see discussion). After correcting for multiple tests using the Benjamini-Hochberg correction, only the correlations between impact load and the relative abundances of *Prevotellaceae,* and *Prevotella* remained significant. Orthopedic injury, caffeine use, pre-workout energy drink consumption, sleep duration, sleep quality, and changes in living or dining circumstances were also significantly correlated with changes in one or more of the four previously mentioned taxa.

**Table 2.** Changes in relative abundances of Coriobacteriales, Prevotella, Prevotellaceae, Ruminococcus, and Verrucomicrobiales correlate with head impacts sustained in the prior 48-72 hours. Dark blue: p < 0.05; light blue: p < 0.10. “Estimate” indicates direction and magnitude of effect on the relative abundance of each taxa.

| Variable | Coriobacteriales |  | Prevotellaceae |  | Prevotella |  | Ruminococcus |  | Verrucomicrobiales |  |
| --- | --- | --- | --- | --- | --- | --- | --- | --- | --- | --- |
|  | Estimate | p (adjusted p) | Estimate | p (adjusted p) | Estimate | p (adjusted p) | Estimate | p (adjusted p) | Estimate | p (adjusted p) |
| Impact Load | -0.0000460 | 0.0154 (0.1593) | -0.0000389 | 0.0045 (0.0492) | -0.0000340 | 0.0035 (0.0437) | 0.0000477 | 0.0075 (0.0773) | 0.0000266 | 0.0775 (0.4803) |
| Time (hour) | 0.0000014 | 0.3851 (0.7598) | 0.0000000 | 0.9983 (0.9983) | -0.0000001 | 0.9303 (0.9581) | 0.0000008 | 0.6000 (0.6889) | 0.0000002 | 0.8807 (0.9973) |
| Impact load*time | 0.0000000 | 0.0111 (0.1593) | 0.0000000 | 0.0064 (0.0492) | 0.0000000 | 0.0056 (0.0437) | 0.0000000 | 0.0181 (0.1121) | 0.0000000 | 0.0619 (0.4798) |
| Player load | 0.0000036 | 0.4167 (0.7598) | 0.0000007 | 0.8172 (0.9983) | 0.0000012 | 0.6677 (0.9581) | -0.0000079 | 0.0623 (0.2452) | 0.0000044 | 0.2230 (0.8640) |
| Orthopedic injury | -0.0039396 | 0.0817 (0.4573) | 0.0005237 | 0.7470 (0.9983) | 0.0002645 | 0.8477 (0.9581) | 0.0006790 | 0.7480 (0.8282) | 0.0042369 | 0.0192 (0.2976) |
| Type 1 - Severe constipation | 0.0091888 | 0.2285 (0.5449) | 0.0000759 | 0.9889 (0.9983) | 0.0002597 | 0.9553 (0.9581) | -0.0107965 | 0.1305 (0.3593) | 0.0022564 | 0.7095 (0.9973) |
| Type 2 - Minor constipation | 0.0047442 | 0.0968 (0.4573) | -0.0001591 | 0.9380 (0.9983) | 0.0002849 | 0.8696 (0.9581) | 0.0018033 | 0.4987 (0.6184) | -0.0019023 | 0.4015 (0.9888) |
| Type 3 - Normal | 0.0025839 | 0.1466 (0.5048) | 0.0001969 | 0.8773 (0.9983) | 0.0002200 | 0.8389 (0.9581) | 0.0023747 | 0.1540 (0.3672) | -0.0006519 | 0.6446 (0.9973) |
| Type 6 - Minor diarrhea | 0.0039896 | 0.1033 (0.4573) | 0.0032928 | 0.0621 (0.3208) | 0.0022477 | 0.1326 (0.5139) | 0.0014749 | 0.5193 (0.6191) | -0.0010788 | 0.5791 (0.9973) |
| Type 7 - Severe Diarrhea | -0.0052824 | 0.4977 (0.8572) | -0.0019435 | 0.7300 (0.9983) | -0.0021526 | 0.6523 (0.9581) | 0.0009056 | 0.9016 (0.9317) | -0.0000396 | 0.9949 (0.9973) |
| Less caffeine | 0.0007833 | 0.8149 (0.9025) | 0.0041892 | 0.0930 (0.4117) | 0.0045349 | 0.0324 (0.1676) | -0.0043551 | 0.1775 (0.3931) | 0.0000092 | 0.9973 (0.9973) |
| Normal caffeine | 0.0001664 | 0.9493 (0.9493) | 0.0019512 | 0.3249 (0.8392) | 0.0020158 | 0.2308 (0.6504) | -0.0018954 | 0.4597 (0.6084) | 0.0001893 | 0.9299 (0.9973) |
| More caffeine | -0.0034689 | 0.6367 (0.8972) | 0.0025396 | 0.6343 (0.9983) | 0.0027741 | 0.5402 (0.9581) | 0.0060439 | 0.3843 (0.5957) | 0.0006844 | 0.9074 (0.9973) |
| Less NSAIDs | -0.0063200 | 0.1824 (0.5141) | 0.0001646 | 0.9615 (0.9983) | 0.0004973 | 0.8634 (0.9581) | 0.0003773 | 0.9322 (0.9322) | 0.0011088 | 0.7687 (0.9973) |
| Normal NSAIDs | -0.0015982 | 0.3947 (0.7598) | -0.0007084 | 0.6021 (0.9983) | -0.0003115 | 0.7869 (0.9581) | 0.0017857 | 0.3128 (0.5335) | -0.0014300 | 0.3410 (0.9888) |
| More NSAIDs | -0.0025527 | 0.6303 (0.8972) | -0.0030882 | 0.4190 (0.8659) | -0.0024684 | 0.4463 (0.9581) | 0.0060080 | 0.2277 (0.4705) | -0.0024832 | 0.5569 (0.9973) |
| Less nicotine | -0.0007253 | 0.8395 (0.9025) | -0.0039514 | 0.1524 (0.5907) | -0.0032316 | 0.1676 (0.5196) | -0.0058522 | 0.1007 (0.3470) | -0.0027836 | 0.3494 (0.9888) |
| Normal nicotine | -0.0011438 | 0.7330 (0.9025) | -0.0022339 | 0.4074 (0.8659) | -0.0022780 | 0.3195 (0.8254) | -0.0054911 | 0.1134 (0.3515) | -0.0005623 | 0.8442 (0.9973) |
| More nicotine | -0.0089504 | 0.2166 (0.5449) | -0.0007383 | 0.8872 (0.9983) | -0.0009373 | 0.8319 (0.9581) | -0.0048889 | 0.4710 (0.6084) | -0.0043883 | 0.4466 (0.9888) |
| Pre-workout drink | -0.0005664 | 0.8443 (0.9025) | 0.0096406 | 0.0000 (0.0002) | 0.0075239 | 0.0000 (0.0011) | 0.0068482 | 0.0129 (0.1003) | -0.0000404 | 0.9861 (0.9973) |
| Number of diet sodas | -0.0013195 | 0.5868 (0.8972) | -0.0000037 | 0.9983 (0.9983) | 0.0000779 | 0.9581 (0.9581) | -0.0005125 | 0.8217 (0.8783) | -0.0002567 | 0.8943 (0.9973) |
| Alcohol | 0.0006860 | 0.0832 (0.4573) | 0.0000280 | 0.9212 (0.9983) | 0.0000778 | 0.7461 (0.9581) | 0.0003034 | 0.4112 (0.5991) | 0.0002611 | 0.4057 (0.9888) |
| Hours of sleep | -0.0017551 | 0.1254 (0.4858) | 0.0018775 | 0.0275 (0.1702) | 0.0016960 | 0.0190 (0.1175) | -0.0033728 | 0.0024 (0.0370) | 0.0007538 | 0.4162 (0.9888) |
| Sleep quality | -0.0004194 | 0.7443 (0.9025) | -0.0010265 | 0.2762 (0.7783) | -0.0005213 | 0.5140 (0.9581) | 0.0014295 | 0.2433 (0.4713) | -0.0021183 | 0.0419 (0.4329) |
| Stress rating | -0.0006042 | 0.1811 (0.5141) | -0.0004269 | 0.2049 (0.7057) | -0.0005154 | 0.0718 (0.3178) | -0.0003475 | 0.4252 (0.5991) | -0.0005337 | 0.1470 (0.6511) |
| Symptom score | 0.0006477 | 0.5566 (0.8972) | -0.0002566 | 0.7456 (0.9983) | -0.0000977 | 0.8841 (0.9581) | -0.0015274 | 0.1391 (0.3593) | 0.0000389 | 0.9646 (0.9973) |
| Minor illness | -0.0002478 | 0.9446 (0.9493) | -0.0029598 | 0.2511 (0.7783) | -0.0030570 | 0.1626 (0.5196) | 0.0033152 | 0.3230 (0.5335) | -0.0011906 | 0.6757 (0.9973) |
| Mild illness | 0.0019500 | 0.6812 (0.9025) | -0.0022947 | 0.5061 (0.9805) | 0.0006965 | 0.8118 (0.9581) | -0.0044012 | 0.3270 (0.5335) | -0.0056542 | 0.1382 (0.6511) |
| Vomitted | -0.0060515 | 0.2909 (0.6441) | -0.0037676 | 0.3591 (0.8564) | -0.0031496 | 0.3661 (0.8730) | -0.0099731 | 0.0633 (0.2452) | 0.0024249 | 0.5941 (0.9973) |
| Change in living or dining | 0.0007368 | 0.8035 (0.9025) | -0.0059301 | 0.0057 (0.0492) | -0.0050722 | 0.0053 (0.0437) | -0.0056016 | 0.0440 (0.2275) | 0.0097826 | 0.0000 (0.0015) |

### Bray-Curtis Dissimilarity increases across the sample collection period

To investigate whether the composition of the gut microbiome varied across the sample collection period, we examined changes in microbial diversity over time (see Methods and Supplementary Figure 2). Comparing early, middle, and late sample collections, we observe an increase in beta diversity as the season progresses (Figure 4a). Friedman’s Chi-Squared test on the Bray-Curtis dissimilarity of the three groups revealed statistically significant differences (χ^2^= 13.44, p = 0.0012, df = 13.44) (Figure 4a). Nemenyi’s post hoc pairwise test revealed that Bray-Curtis dissimilarity was significantly lower early in the collection period (0.280 ± 0.072) than late (0.326 ± 0.048, Q_34_ = 3.71, p = 0.001).

**Figure 4.**
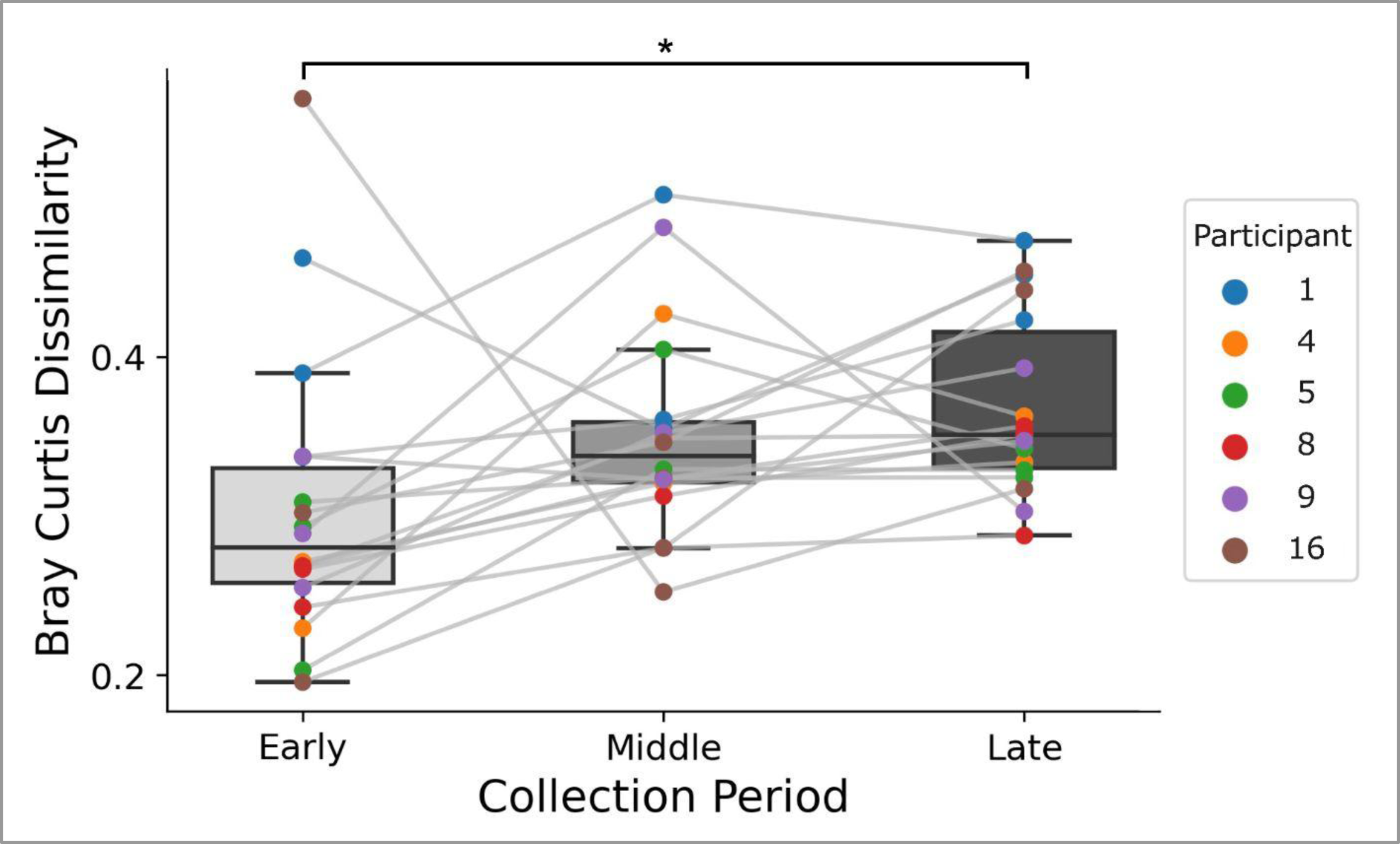
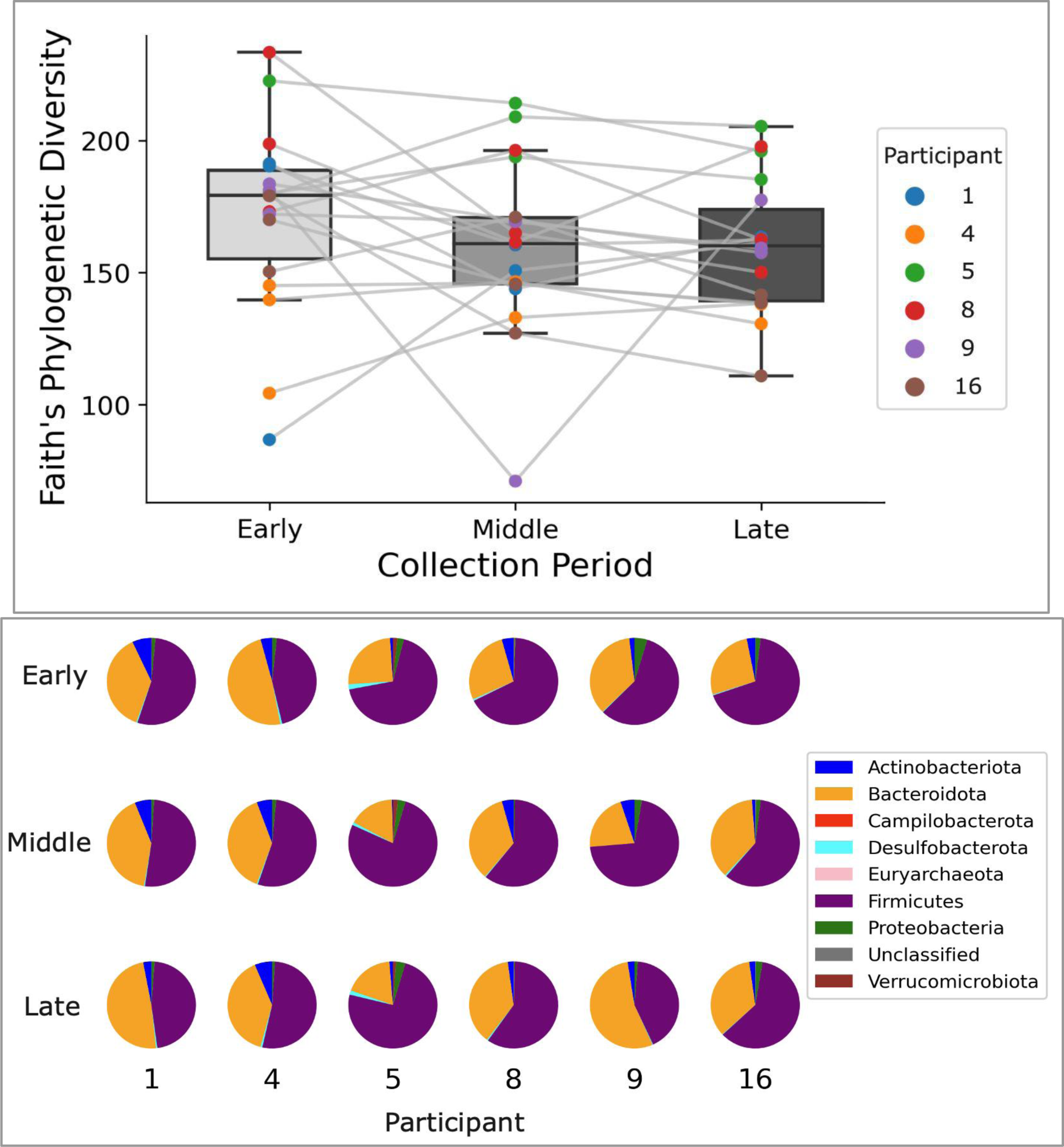
(a) Bray-Curtis Dissimilarity increases across the sample collection period. The first three, the center three, and the last three data points available for each participant (excluding postseason samples) were used for the early, middle, and late periods, respectively (see methods and Supplementary Figure 2). Bray-Curtis Dissimilarity increased from the early to late period (*indicates p < 0.05). **(b)** *Faith’s Phylogenetic Diversity did not change significantly across the sample collection period*. Analysis revealed no significant differences in Faith’s Phylogenetic Diversity across the sample collection period for the three timepoints analyzed. **(c)** *Relative abundance of phyla varies across participants and across the sample collection period.* Firmicutes and Bacteriodota were the most abundant phyla for all participants, which varied among participants and across the sample collection period.

We also investigated changes in alpha diversity across the sample collection period as measured by Faith’s Phylogenetic Diversity (Figure 4b). Friedman’s Chi-Squared test on Faith’s Phylogenetic Diversity revealed no significant differences between any time periods (χ^2^ = 3.0, p = 0.2231). While Firmicutes and Bacteroidetes were the most abundant phyla for all participants, their relative abundance varies (Figure 4c). Some variation does occur within individuals across the collection period, with fluctuations in the abundant phyla Firmicutes and Bacteroidetes most visible for players 4, 5, and 9.

## DISCUSSION

The present study aimed to determine if and how the human gut microbiome changes in response to SHIs. The rationale for the present study is based on previous literature that demonstrates changes in the gut microbiome following mTBI in murine models and the limited available evidence demonstrating similar findings in humans^40–49^. Amongst the six NCAA Division I Football players who participated, our results provide evidence that SHIs sustained in practices and competitions led to alterations in the gut microbiome.

The six participants sustained an average of 261 (SEM 51.9) head impacts and an average impact load score of 6,308.25 (SEM 1,435.2) across the season (52.0 (SEM 11.8) impact load sustained per practice session and 260.0 (SEM 86.9) impact load sustained per game session) (Figure 2b). This number of impacts sustained across the season and per session was consistent with other studies using the same head impact monitoring system on collegiate football players^61^. Isolating incidents in which there was substantial head impact exposure followed by multiple days of subthreshold head impact exposure revealed that Bray-Curtis Dissimilarity increases with 48 to 72 hours of head impact exposure, which was confirmed with mixed-effect linear models accounting for 15 clinical factors. These models also demonstrated that decreases in the relative abundances of Coriobacteriales*, Prevotellaceae,* and *Prevotella* and increases in *Ruminococcus* and *Verrucomicrobiales* were correlated with greater head impact exposure. The acute increases in Bray-Curtis dissimilarity recognized following SHIs and the longitudinal increase in dissimilarity we identified across the competition season suggest that the accumulation of SHIs potentially contributes to chronic, longitudinal changes in gut microbial diversity.

Repeated measures ANOVA and mixed-effect linear models demonstrated that increases in Bray-Curtis Dissimilarity were correlated with SHI load 48 to 72 hours after exposure. A previous study in rats demonstrated that the gut microbiome was stable for at least 24 hours following head impact, while significant changes in microbial diversity were observed 72 hours following mTBI^49^. Given this overlap in the onset of microbiome changes between this study and previous studies, it is likely that SHIs and mTBIs may take between 48 and 72 hours to lead to changes in gut microbial diversity. Previous works have reported that cognitive deficits such as memory impairment, decreased executive function and response time, as well as increases in peripheral neuroinflammatory markers, including S100 calcium-binding protein beta (S100B) and neurofilament light (NfL)^67,86–89^, appear to occur before the shift in microbial diversity others have observed in rodent models of mTBI and that we report here^48^. Thus, a short-term analysis examining fluctuations in peripheral biomarkers of neuroinflammation and cognitive function following SHI and mTBI could determine if the acute changes in the gut microbiome following brain trauma are linked with the severity of cognitive deficits and pro-inflammatory profiles.

Comparing the alterations in the gut microbiome between human subjects is challenging because it has been demonstrated that individual gut microbiomes can have unique responses to the same input, likely caused by the combined effects of unique host genetics and current microbial composition^90,91^. Nonetheless, our data suggest that the alterations in the relative abundances of specific taxa that we identified match those previously identified in the literature examining brain injury and the microbiome and are functionally relevant.

We identified that decreases in the relative abundance of the microbial order Coriobacteriales were correlated with head impact loads sustained in the past 48-72 hours. To our knowledge, Coriobacteriales has not been identified as changing in response to brain injury in murine and human models (Supplementary File 3). However, a decreased relative abundance of Coriobacteriales has been associated with mild cognitive impairment in hypertensive patients, known to be at a higher risk for dementia^92^.

Similar to our results, the microbial family *Prevotellaceae* has been found to decrease in response to TBI in mice, rats, and humans (with the change retained years later, in the case of humans)^47,77^. Depletion of *Prevotellaceae* has been associated with increased production of pro-inflammatory cytokines TNF-α and IL-1β in serum and decreased butyrate production in the gut, which has been shown to provide a neuroprotective effect by restoring the blood-brain barrier (BBB) following TBI.^93,94^ Additionally, the *Prevotella* genus has been demonstrated to increase the concentration of short-chain fatty acids (SCFAs) in pigs; SFCAs inhibit inflammation and are potentially protective against dementia risk^95,96^. Yet, certain species of *Prevotella* (a genus within *Prevotellaceae*) have been shown to decrease SCFA production and perpetuate intestinal inflammation^97^. Thus, the effect of depletion in *Prevotellaceae* and *Prevotella* likely depends on the host’s microbial composition and the specific species present.

Moreover, we found that sustaining a head impact is correlated with an increase in the *Ruminococcus* genus 48-72 hours following the impact; however, in response to brain injuries, other studies have found decreases in the *Ruminococcaceae* family and a species of *Ruminococcus* in murine models but long-term increases in the *Ruminococcaceae* family in humans with severe TBI^46–48,79^. The *Ruminococcus* genus and species within the genus are increased in patients with inflammatory bowel disease and have been associated with producing pro-inflammatory metabolites in Crohn’s disease^98,99^.

We also found a marginally significant association between head impact load and increases in *Verrucomicrobiales*. *Listeriaceae* is a family member within this order, which increases in abundance in the gut in response to mTBI in rats^48^, and the phylum *Verrucomicrobia* has been reported to increase in response to TBI in humans^47^. The family *Verrucomicrobiaceae* also increases in response to acute inflammation in mice, and the phylum *Verrucomicrobia* was shown to induce inflammation in the colon of rats^100,101^.

The changes in the relative abundance of these taxa with SHIs and the relevant relationships between the taxonomic shifts and inflammatory states suggest that SHIs may nudge the gut microbiome towards an inflammation-promoting state that could contribute to longer-term neurological consequences. However, the inconsistency of the trends in taxonomic changes across the literature and between organisms indicates that future explorations across a more diverse population are necessary to determine if these taxonomic changes following exposure to SHIs are consistent. To find out if these changes in taxonomy are linked to inflammatory traits, these studies might also measure the amounts of SCFA in stool and levels of serum biomarkers before and after being exposed to SHIs.

The mathematical modeling of our data provides further evidence for a link between SHIs and changes in microbiome composition. Notably, the interaction effect between head impact load and time (“impact load*time”) was significant in the models for all of the aforementioned taxa (Table 2). The interaction effect may point to changes in response to head impacts that depend on time or host (note the uneven distribution of sample collections between participants over time; see Figure 2a, Supplementary Figure 2). This would suggest that in-depth, longitudinal analyses of individual microbiomes may be more informative when attempting to determine the relationships between certain factors and specific taxa compared to group-wide or population studies, particularly when studying humans with distinct microbial profiles.

Furthermore, the mixed-effect linear model demonstrated correlations between time (across the entire study period), player load (a measure of physical activity during each practice and game), NSAID use, and consumption of pre-workout energy drinks and increasing dissimilarity in gut microbial composition from the start of the season. The correlation between time and increasing microbiome dissimilarity between players might be explained by other lifestyle changes associated with athletic competition seasons that were not accounted for^90^. Similarly, exercise intensity, NSAID use, and ingredients commonly found in pre-workout supplements have been shown to change the diversity and composition of the gut microbiome in humans^102–104^. Reporting “less than normal” and “more than normal” NSAID use was not significantly correlated with Bray-Curtis Dissimilarity (Table 1), which was likely the case because participants sparsely reported this behavior (Supplementary Table 1).

Higher stress ratings were associated with decreasing Bray-Curtis dissimilarity compared to the baseline samples, likely due to the higher reported levels of stress at the beginning of preseason training camp within this study and generally across collegiate athletes^105^. Given that collegiate athletes partake in cyclical behaviors due to the weekly practice and competition cycles, further investigation might include an in-depth analysis of dietary behaviors, activity patterns (outside of those for the sport), and other behaviors to confirm that the associations between SHIs and gut microbiome changes are not due to another confounding factor. Moreover, an analysis with additional dependent variables, such as serum inflammatory cytokines and cognitive assessments, might identify other factors outside of brain injury contributing to chronic inflammation and long-term cognitive deficits among American football players.

Although we observed five taxa that underwent an acute response to SHIs, we did not identify significant changes in the relative abundances of any of these taxa across the collection period. Additionally, we did not detect significant changes in alpha diversity across this period (Figure 4b). The small cohort size and the relatively light impact load burden that Participants 4, 8, and 16 experienced throughout the season could contribute to this lack of effect. However, a visual downward trend in alpha diversity is visible across the collection period, suggesting a trend toward dysbiosis (Figure 4b). A larger sample size is needed to confirm this downward trend and its potential association with SHIs sustained across the season.

An increase in Bray-Curtis Dissimilarity was observed across the season (Figure 4a), which, due to the significant association between Bray-Curtis Dissimilarity and head impact load (Table 1), suggests a potential cumulative effect of SHIs leading to long-term alterations in the gut microbiome. A larger cohort is needed to confirm and characterize this effect. However, the accumulation of SHIs leading to long-term alterations in the gut microbiome could have implications for the chronic inflammation associated with repeated mTBI, which may be linked to short-term disability, neurodegeneration, and early cognitive decline^51–53^.

This pilot study provides the first evidence that subconcussive head impacts lead to changes in the diversity and composition of the gut microbiome. Potentially pro-inflammatory-related changes in four microbial taxa were correlated with SHI load, and longitudinal changes in microbial diversity across the competition season suggest that SHIs may nudge the human gut microbiome towards a state that promotes chronic inflammation and, thus, an increased risk for neurodegeneration. Future studies should include a larger and more diverse population that includes females, as they are known to respond differently to mTBIs^106^, monitor more factors, and include peripheral biomarkers and cognitive assessments.

## ACKNOWLEDGEMENTS

This study could not have been accomplished without the Colgate football players who volunteered for study, Dom Calhoun and the Colgate athletics equipment staff, and Bryan Klobucar, Matt Shimshock and colleagues at Riddell. We also thank Cody Herbert, Jillian Austin-Pottorff, and Stan Dakosty and the Colgate Football coaching staff for their input and support, Chuck Monteith, Paul Klawitter, and Dong Wang for intellectual contributions, and Kristen Dams-O’Connor and Verena Link for critical reading of the manuscript. This project was funded by Colgate University via the Russell Colgate Distinguished University Professorship and Stuart Updike Fund for Undergraduate Research.

## AVAILABILITY of DATA and MATERIALS

The code used for data processing and statistical analyses is available on this GitHub repository: https://github.com/aziz-zafar/TBI-Microbiome

Sequence data used in this study have been deposited with the National Center for Biotechnology Information as BioProject ID PRJNA1111907. Please contact KDB with questions or requests for additional information.

