## Supplemental tables and figures for "Subconcussive head impacts sustained during American football alter gut microbiome diversity and composition"

### SUPPLEMENTARY MATERIALS

| | Factors | Description or rationale for exclusion | Mean and SEM or frequency |
|---|---|---|---|
| *Sport-related factors* | Impact load | calculated for each session based number of impacts and impact intensity | 66.6 ± 9.82 |
| | Player load | measure of effort exerted during each session from Catapult Vector | 1170 ± 42.4 |
| | Orthopedic injury | currently being treated by athletic trainer for injury (binary) | yes = 31; no = 195 |
| *Fecal sample information* | Time | time of fecal sample collection (total hour of study) | - |
| | Bristol Stool Form Scale | Bristol Stool Chart (Type 1- Type 7) | 1 = 3 ; 2 = 17 ; 3 = 61; 4 = 117; 5 = 0; 6 = 2; 7 = 2 |
| *Drugs & diet information* | Caffeine use | (1) none at all, (2) less than normal, (3), normal, (4) more than normal | 1 = 70; 2 = 24; 3 = 128; 4 = 4 |
| | NSAID use | (1) none at all, (2) less than normal, (3), normal, (4) more than normal | 1 = 71; 2 = 6; 3 = 144; 4 = 5 |
| | Nicotine use | (1) none at all, (2) less than normal, (3), normal, (4) more than normal | 1 = 157; 2 = 20 ; 3 = 46; 4 = 3 |
| | Preworkout | binary | yes = 24; no = 202 |
| | Diet sodas | number of diet sodas | 0.0885 ± 0.0209 |
| | Alcohol | number of standard drinks | 0.553 ± 0.139 |
| *Health information* | Hours of sleep | total hours of sleep (max = 10) | 7.430 ± 0.0584 |
| | Sleep quality | (0) very poor - (4) very good | 2.84 ± 0.0597 |
| | Stress rating | five questions adopted from PSS-10 (min = 0; max = 20) | 2.04 ± 0.140 |
| | Symptom score | five questions adopted from SCAT-5 (min = 0; max = 20) | 0.230 ± 0.0474 |
| | Illness | (1) none, (2) minor, (3) mild, (4) severe | 1 = 209; 2 = 11; 3 = 6; 4 = 0 |
| | Vomitting | binary | yes = 4; no = 222 |
| *Other* | Change in living or dining circumstance | changing dorms or meal plans | twice for each participant |
| *Excluded Factors* | Stool color | few reported cases of anything except "brown" | - |
| | Blood in stool | only one reported case | - |
| | Fiber consumption | homogenous responses; error in reporting methods | - |
| | Red meat consumptoion | homogenous responses; error in reporting methods | - |
| | Carbohydrate consumption | homogenous responses; error in reporting methods | - |
| | Protein powder | only used by one participant | - |
| | Multi-vitamin | only used by one participant | - |
| | Fish oil | only used by one participant | - |
| | Prescription medications | only used by one participant | - |
| | Cannabis use | not heavily used; participant confidentiality | - |
| | Suspected head impact on the field | use to verify head impact montioring | - |
| | Suspected head impact off the field | none reported | - |

**Supplementary Table 1**

*Clinical factor descriptions, responses, and exclusions.* 30 total factors were monitored across the duration of the study; 28 were acquired from survey data, and 2 were acquired from on-field devices. 12 factors acquired from survey data were excluded due to low reporting or errors in reporting methods. Three dietary factors were excluded due to the flows in the collection methods and the homogeneity of responses. See Supplementary Files 1 and 2 for survey questions and formats.

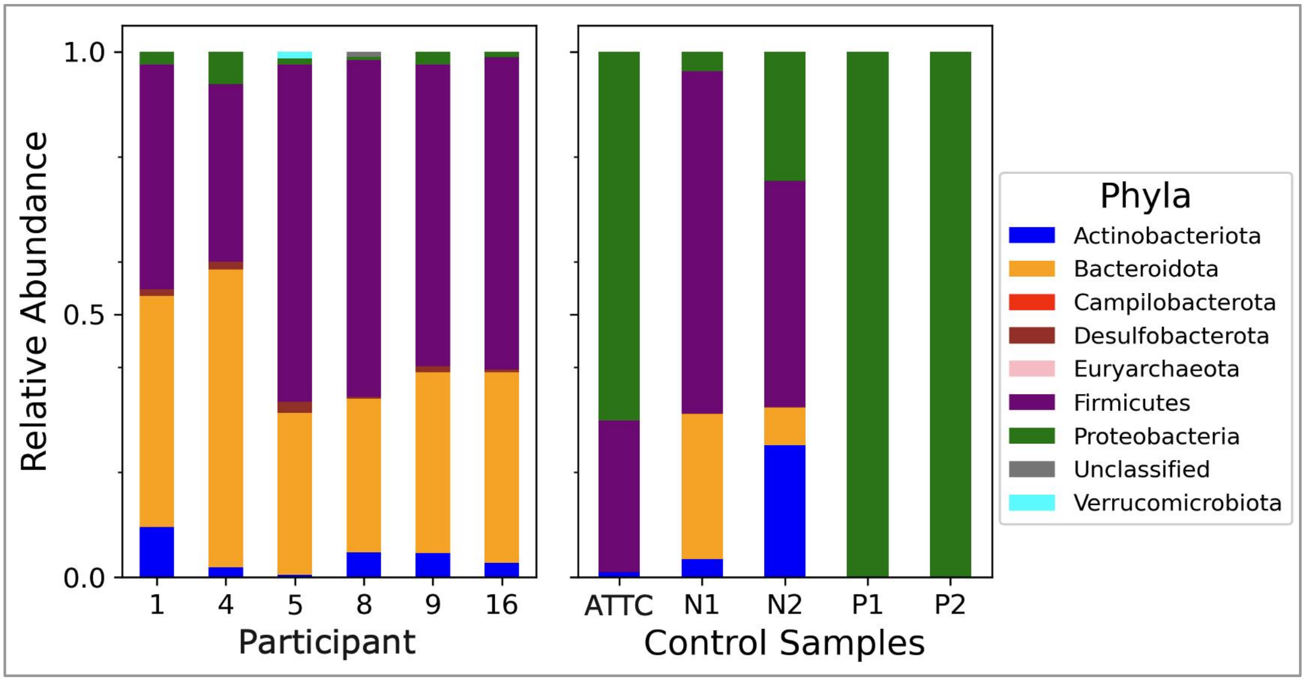


**Supplementary Figure 1**

*Analysis of control samples.* 16S amplification and DNA sequence analysis were performed on five non-fecal samples as controls for taxonomic analysis. The left panel shows the distribution of phyla in the preseason baseline sample from each of the six participants. The right panel shows the distribution of phyla across the five control samples. The internal mock from ATCC, is the ABRF-MGRG 10-strain even mix MSA-3001 (ATCC, Gaithersburg, MD, USA). N1 was a blank sample that went through the DNA extraction process, and N2 was the stock solution that the DNA was dissolved in. Positive controls (P1 and P2) consisted of 100% Escherichia coli.

| Characteristics | Participant | | | | | | | |
|---|---|---|---|---|---|---|---|---|
| | 1 | 4 | 5 | 8 | 9 | 16 | Avg | SEM |
| Height (cm) | 190.5 | 185.4 | 177.8 | 193.0 | 188.0 | 188.0 | 187.1 | 2.1 |
| Body weight (kg) | 138.3 | 97.5 | 83.9 | 108.9 | 104.3 | 111.1 | 107.4 | 7.4 |
| BMI | 38.1 | 28.4 | 26.5 | 29.2 | 29.5 | 31.4 | 30.5 | 1.7 |
| Days since concussion | 1,809 | N/A | 299 | 284 | 341 | 1,572 | - | - |
| Days since oral antibiotic | 64 | 99 | N/A | N/A | N/A | N/A | - | - |
| Head impacts sustained | 348 | 176 | 451 | 203 | 324 | 64 | 261 | 56.9 |
| Head impact load sustained | 7,139.0 | 4,714.5 | 12,709.5 | 4,482.0 | 7,512.0 | 1,292.5 | 6,308.3 | 1572.1 |
| Head impact load per practice | 54.8 | 20.0 | 118.1 | 30.0 | 73.2 | 16.1 | 52.0 | 15.9 |
| Head impact load per practice SEM | 10.2 | 5.3 | 18.8 | 6.2 | 13.1 | 6.2 | 11.8 | - |
| Head impact load per game | 320.4 | 307.0 | 445.2 | 227.6 | 239.2 | 20.6 | 260.0 | 57.4 |
| Head impact load per game SEM | 129.0 | 73.4 | 100.3 | 63.1 | 54.9 | 11.4 | 86.9 | - |
| Fecal samples analyzed | 83 | 39 | 20 | 27 | 31 | 26 | 37.7 | 9.4 |
| Lifestyle questionnaires completed | 82 | 20 | 21 | 25 | 29 | 23 | 33.3 | 9.8 |

**Supplementary Table 2**

*Baseline participant demographics, medical history, sports information, and data availability.* Days since the last concussion and oral antibiotic use are calculated from the first day of fecal sample collection. The total number of head impacts and impact load sustained were measured throughout the entire season that the study was conducted.

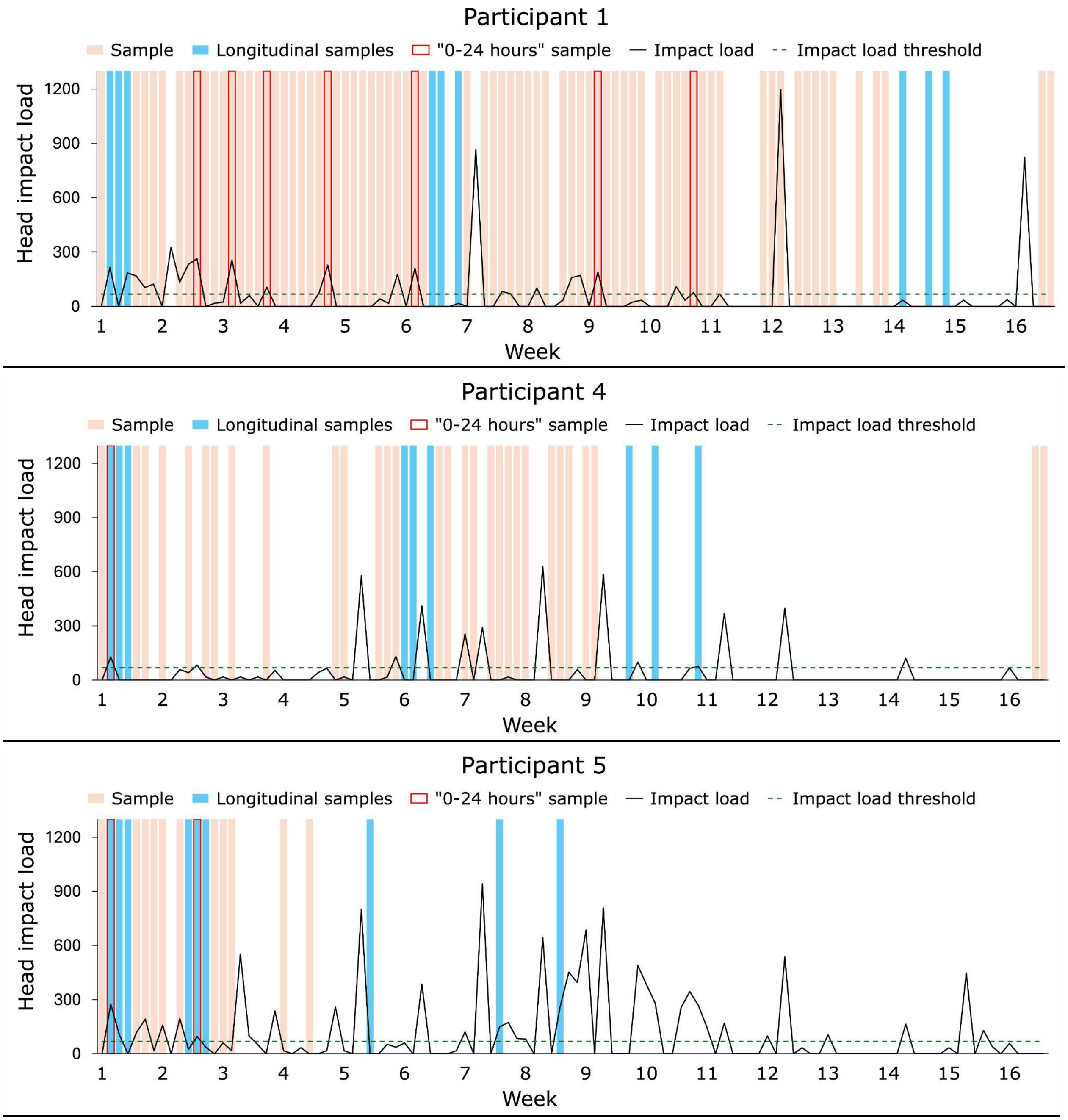
Participant 1
Sample
Longitudinal samples
"0-24 hours" sample
Impact load
Impact load threshold
Head impact load
1200
900
600
300
0
1 2 3 4 5 6 7 8 9 10 11 12 13 14 15 16
Week
Participant 4
Sample
Longitudinal samples
"0-24 hours" sample
Impact load
Impact load threshold
Head impact load
1200
900
600
300
0
1 2 3 4 5 6 7 8 9 10 11 12 13 14 15 16
Week
Participant 5
Sample
Longitudinal samples
"0-24 hours" sample
Impact load
Impact load threshold
Head impact load
1200
900
600
300
0
1 2 3 4 5 6 7 8 9 10 11 12 13 14 15 16
Week

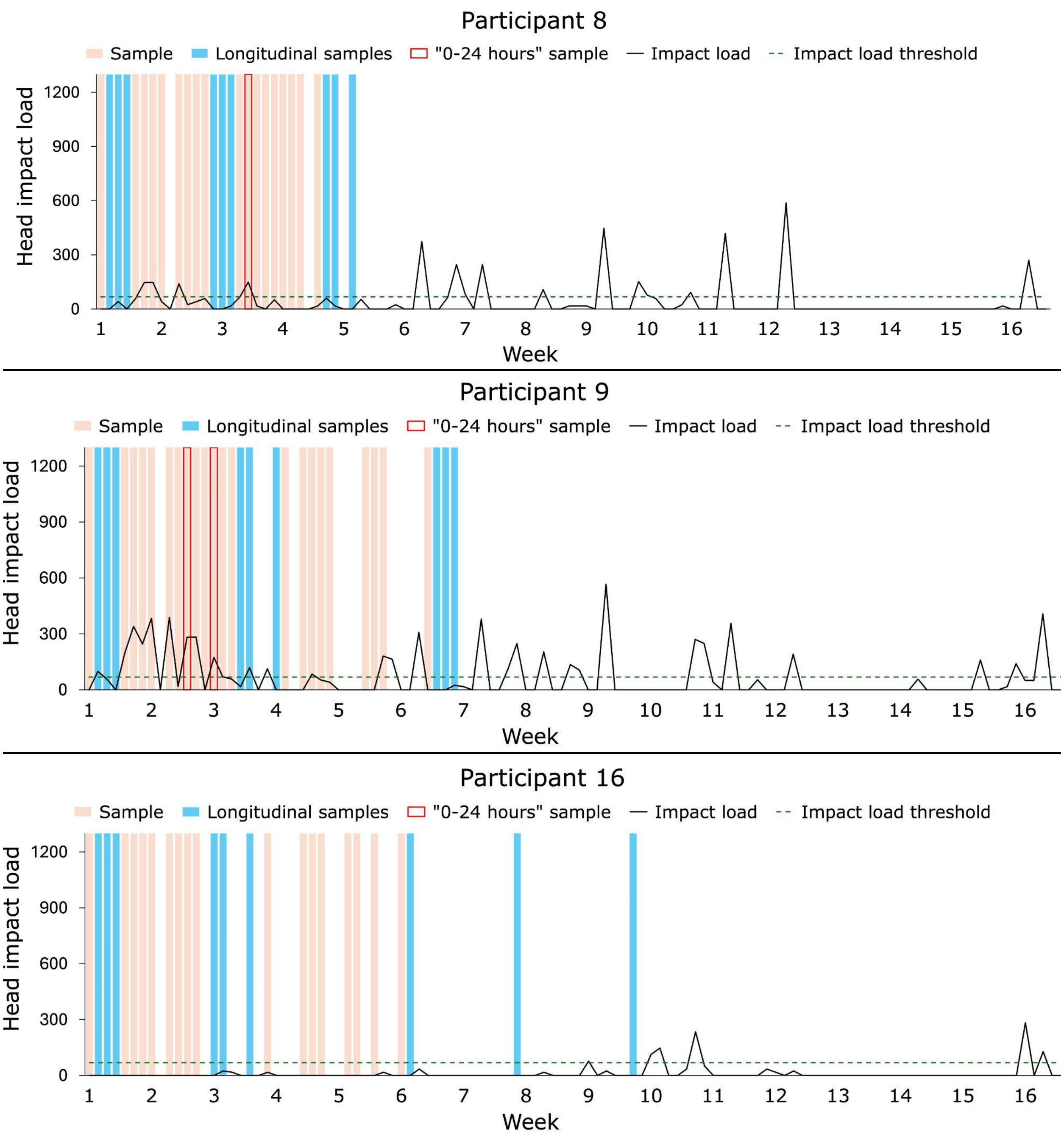


**Supplementary Figure 2**

*Sample collection and head impact overview.* Each vertical bar represents an individual date across the collection period. The days on which samples were collected are colored orange or blue. The samples used in the longitudinal analysis are colored blue (see methods and Figure 4). The dashed green line indicates the threshold for “substantial head impact exposure” (see methods). Samples outlined in red indicate the starting point (0-24 hours post substantial head impact exposure) for the analysis in Figure 3 (see methods).

**Supplementary File 1**. *Participant background questionnaire.* The survey was completed through Google Forms.

**Supplementary File 2**. *Fecal sample collection lifestyle questionnaires.* The survey was completed through Google Forms.

**Supplementary File 3.** *Literature review of the taxa used in analysis*. The taxa used in analysis are highlighted in orange.

**Supplementary File 4.** *All mixed-effect linear models.*

### CODE AVAILABILITY

The code used for data processing and statistical analyses is available on this GitHub repository: https://github.com/aziz-zafar/TBI-Microbiome

### SEQUENCE DATA AVAILABILITY:

Sequence data used in this study have been deposited with the National Center for Biotechnology Information as BioProject ID PRJNA1111907. Please contact KDB with questions or requests for additional information.
